# Lesions of canine prostate underwent laser ablation, radiofrequency ablation, and microwave ablation: Lesions’ shapes and clinical significances

**DOI:** 10.1101/774455

**Authors:** Liu Ruiqing, Zhang Lianzhong, Duan Shaobo, Cao Huicun, Cao Guangshao, Chang Zhiyang, Zhang Ye, Li Yaqiong, Wu Yuejin, Liu Luwen

## Abstract

**Objective:** To explore the shapes and clinical significances of the lesions of canine prostate underwent laser ablation (LA), radiofrequency ablation (RFA), and microwave ablation (MWA).

**Methods:** Here, 6 male Beagle dogs were randomly assigned to LA, RFA, and MWA groups, respectively. The ablations were conducted with the doses commonly applied in clinical practice (LA: 3 W/1200 J; RFA and MWA: 30 W/120 s) for one region in the left and right lobes of the prostate, which totally resulted in 12 lesions. An energy transmitter was inserted via the perineum under the guidance of transrectal ultrasound (TRUS) and the ablation process was observed dynamically. After ablation, the efficacy was assessed by contrast-enhanced ultrasonography (CEUS), and transverse diameter (TRD), anteroposterior diameter (APD), and longitudinal diameter (TRD) were measured. Then, the volume (V) was calculated according to the following formulae, V = 1/6 × π × (MLD × APD × LD), and ratio (R) R =[(MLD + APD)/2 × LD], which was used to show the shape of the lesion (The closer R is to 1, the more spherical it is).

**Results:** The shapes and sizes of the lesions underwent the three techniques varied under the clinically commonly used doses. The R values, reflecting the changes in shapes of lesions, were 0.89±0.02, 0.72±0.01, and 0.65±0.03 for RFA, MWA, and LA, respectively, which were significantly different (*P* =0.027). The volumes of the lesions underwent the three ablation techniques significantly varied as well (P=0.001), which were 2.17±0.10, 1.51±0.20, and 0.79±0.07 ml for MWA, LA, and RFA, respectively. Conclusion: The above-mentioned three techniques could be used for ablation of prostate cancer using the clinically commonly used doses. The shapes of the lesions underwent the three ablation techniques varied evidently, with the lesions received RFA and LA were more spherical and oval, respectively, while the shapes of lesions underwent MWA were between spherical and oval. The lesions received MWA and RFA had the largest and smallest sizes, respectively, while sizes of lesions underwent LA were in the middle. These findings may assist physicians to select an appropriate therapeutic method for treatment of prostate disease.

## Introduction

Benign prostatic hyperplasia (BPH) is a disease frequently found in middle-aged and old men, which could induce lower urinary tract symptoms (LUTS), and therefore, severely influence patients’ quality of life [1]. Drug therapies and surgical treatments have been used for the treatment; however, both approaches accompany with a number of side-effects [2]. A study on epidemiology showed that the incidence and mortality of prostate cancer (PCa) are remarkable in males [3]. The conventional therapeutic strategies include operation, radiotherapy, and endocrinotherapy [4], however, these treatments all have certain influences on sexual functions, urination, and defecation [5]. Local minimally invasive treatment possesses several great advantages, including being minimally invasive, repeatable, as well as associating with less side-effects. Therefore, increasing attentions have been paid on this method, which has been extensively applied in clinical practices [6,7]. Laser ablation (LA), radiofrequency ablation (RFA), and microwave ablation (MWA) are thermal ablation techniques, which could induce high temperature within the tissues, and may lead to irreversible damages to tissues. LA has already been conducted in a variety of countries for the treatment of prostate disease [8,9], and the effectiveness and safety profiles have been demonstrated. The FRA and MWA were conducted in animal experiments as well. Pathological examinations have demonstrated that RFA and MWA could induce coagulative necrosis. The high-power MWA (45 W/180 s) results in he penetration of ablation through the prostate, suggesting that appropriate power and time should be taken for the treatment into consideration [10,11]. To date, no studies have reported which of the three ablation techniques should be selected for BPH and PCa. In the present study, animal experiments were conducted to explore the characteristics related to shapes and clinical significances of the lesions of canine prostate underwent the three types of ablation techniques, using the most frequent clinically used doses. Our findings may provide assist physicians to select an appropriate ablation technique for thermal ablation of prostate diseases in clinical practice.

## Materials and Methods

### Animals

Here, 6 male Beagle dogs with mating history were purchased from Dilepu Biomedical Co., Ltd. (Xian, China); The animals were weighed and were randomly assigned to LA, RFA, and MWA groups, respectively . their mean age was 5.2±0.8 years old, and their body weight was 14.2±2.2 kg. The dogs were kept in the Center for Drug Safety Evaluation of Zhengzhou University (Zhengzhou, China). The animals were individually housed in stainless steel cages (L 1000 × W 1000 × H 2100 mm) before and after ablation. The environment in which dogs live is set to 23 ± 3 °C of temperature, 55 ±15 % of relative humidity, 10~20 times/hr of ventilation frequency, 12 h of lightingduration (lighting up at 8 a.m. ~ lighting out at 8 p.m.) and 150~300 Lux of luminous intensity. Each dogs was offered a daily ration of 300 g of solid food (Beijing Keao Xieli Feed Co.,Ltd.). The uneaten food was collected on the following morning. Water was disinfected by ultraviolet sterilizer and ultrafiltration and made available ad libitum using an automatic water supplier.

The study protocol was approved by the Ethics Committee of Center for Drug Safety Evaluation of Zhengzhou University (Zhengzhou, China).

### Equipment

An ultrasound system (BK3000; B-K Medical Systems, Inc., Peabody, MA, USA) equipped with a transrectal probe and contrast tuned imaging (CnTI) were used in the current study. Magnetic resonance imaging (MRI) was conducted by using Signa HDxt 3.0T (GE Healthcare, Chicago, IL, USA).

For the LA ablation, Echolaser X4 (Esaote SpA, Genoa, Italy) with the emission source of Nd:YAG laser (wavelength of 1064 nm) was used, under the guidance of 21G puncture needle. The length of the fibers was 20 cm, which were inserted at 1.0cm through the puncture needle. For RFA, LDRF-120S RFA instrument (Lead Electron Corp., Mianyang, China) and lddjs3-0080100B radiofrequency electrode were used, with the working tip of 1 cm. For MWA, ECO-100E1 (MWA, Ecosystems Ltd., Leigh, UK) with the frequency of 2450 MHz and maximal output power of 100 W was utilized. The outer diameter of the antenna (ECO-100AL2) was 1.4 mm, and the length of the electrode was 15 cm. The instruments for RFA and MWA were equipped with an internal cooling system to ensure that the temperature of the needle trunk was < 10 °C.

### Contrast agent and anesthetics

SonoVue (Bracco Group, Milan, Italy) was used as a contrast agent for ultrasound examinations in the present study. For this purpose, bolus injection of the contrast agent with the dose of 0.04 ml/kg was conducted [12]. Magnevist® with the dose of 0.21 ml/kg was herein used as the contrast agent for MRI.

Pentobarbital sodium (Sinopharm Chemical Reagent Co., Ltd., Shanghai, China) with the dose of 15 mg/kg and Zoletil 50 (Virbac Inc., Carros, France) with the dose of 0.2 ml/kg were used as the anesthetics in this study.

### Methods

#### Preoperative preparation

The dogs were fasted for 12 h, then, the skins of the abdomen and perineum were prepared, and cleansing enema was conducted. An indwelling needle was implanted at the forearm. After that, total intravenous anesthesia (TIVA; intravenous injection of pentobarbital sodium followed by anesthesia maintenance with Zoletil 50) was performed, the dogs were placed in supine position, and the four extremities were fixed. Next, topical disinfection and draping were undertaken.

#### Ablation

The ablation in all the dogs were conducted by a physician with ablation experience of over 5 years. Transrectal ultrasound (TRUS) examination was initially conducted for the gross scanning of prostate. The size of the prostate was measured, and then, the site for puncture was selected with the aid of perineal approach. Afterwards, an energy transmitter was inserted via perineum under the guidance of TRUS. The distance from the tip of the transmitter to urethral canal, prostatic capsule, rectum, and bladder was > 6 mm, while the distance from the tip of the laser fiber to bladder was > 15 mm (Fig. 1). The ablation process was dynamically monitoring by TRUS. For each dog, the ablation was conducted at both the left and right lobes of prostate to generate one lesion, according to the doses used in the thyroid gland that previously reported [13–15]. Eventually, the ablation resulted in 12 lesions in the dogs, with each ablation technique resulted in 4 lesions. The power and energy required for LA were 3 W and 1200 J, while the power and time required for RFA and MWA were 30 W and 120 s.

**Fig 1.**
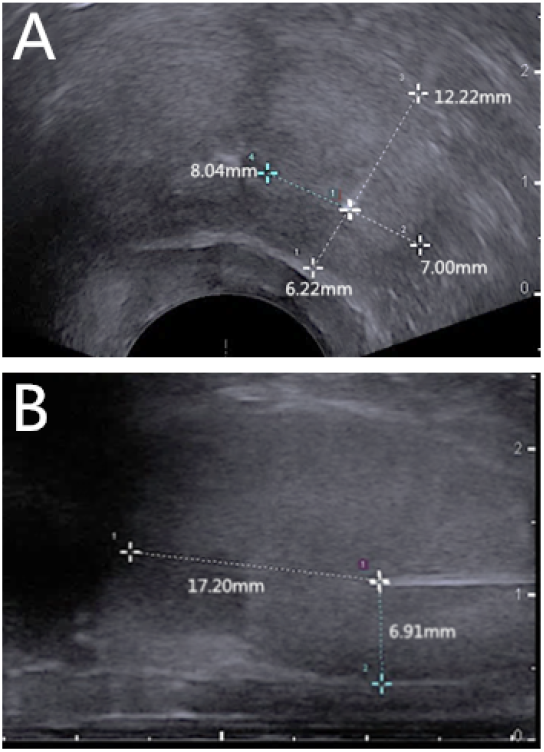
Distances from the tip of the laser fiber to the capsules of different organs on transrectal ultrasound images of prostate (A, transverse section; B, longitudinal section)

#### Postoperative assessment

To minimize the influences of hyperecho images induced by the gas during ablation on imaging effects, transrectal contrast-enhanced ultrasonography (CEUS) was conducted for the prostate of each dog at about 10 min after ablation [12]. In brief, the instrument was set at the imaging mode, bolus injection of SonoVue (0.04 ml/kg) was carried out through the indwelling needle at the forearm, and then, 5 ml normal saline was used to rinse the tube. The CEUS image of each lesion was observed, and transverse diameter (TRD) and anteroposterior diameter (APD) were measured on the transverse section of the lesion, while longitudinal diameter (LD) was measured at the longitudinal section (Fig. 2). The ratio (R), which was used to show the shape of lesion, was calculated according to the following formulae: R = (MLD+ APD)/2LD. The closer R is to 1, the more spherical it is [16,17]. The volume (V) of the lesion was calculated as follows: V=1/6×π×(MLD×APD×LD).

**Fig 2.**
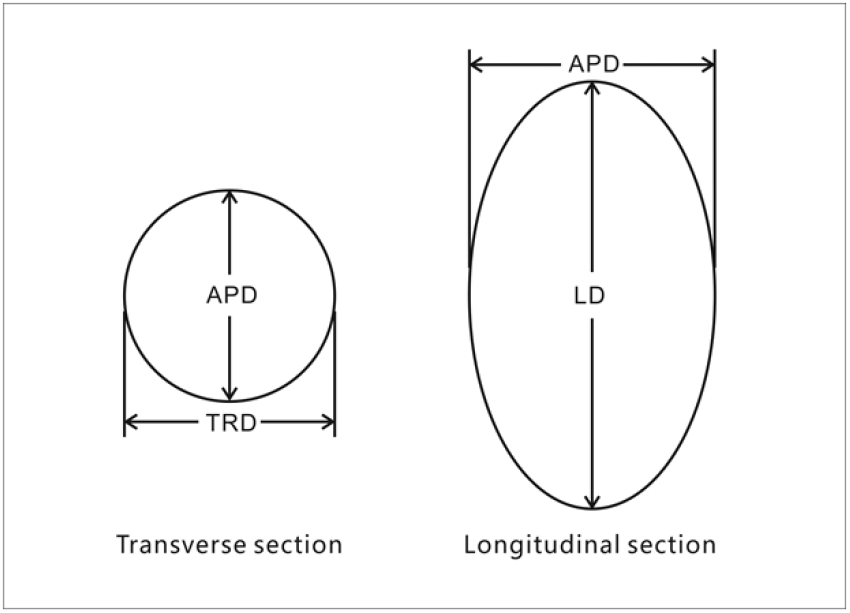
The TRD, APD, and LD measured in the transverse and longitudinal sections.

#### Postoperative care

The breathing and heartbeat were closely monitored for each dog during 6 h after ablation. The dogs were offered food and water after ablation 12 h.To prevent infection, each dog was intramuscularly administered 80mg/kg gentamicin (Sinopharm Chemical Reagent Co., Ltd.) for one week following the end of abltaion. Besides, we pay attention to the symptoms of hematuresis, hematochezia and pain about dogs. After the completion of follow up observation, the animals were sacrificed by injection pentobarbital sodium (30mg/Kg) at end of experiment.

### Statistical analysis

In the present study, SPSS 22.0 software (IBM, Armonk, NY, USA) was used to perform statistical analysis. Quantitative data were described as mean ± standard division (SD). Student’s t-test was utilized to compare size of lesions, while non-parametric rank-sum test was used for making comparison among different groups. P<0.05 was considered statistically significant.

## Results

Two-dimensional TRUS showed that hyperechoic regions were gradually expanded during LA, RFA, and MWA of prostate of dogs, which was followed by acoustic shadow or acoustic tail. The acoustic shadow disappeared after the ablation was completed. Besides, CEUS showed that the lesions were spherical (transverse section, Fig. 3A) or oval (longitudinal section, Fig. 3B), which were without contrast agent filling; the boundaries of the lesions were clear, and the shapes of the lesions were regular (Fig. 3).

**Fig 3.**
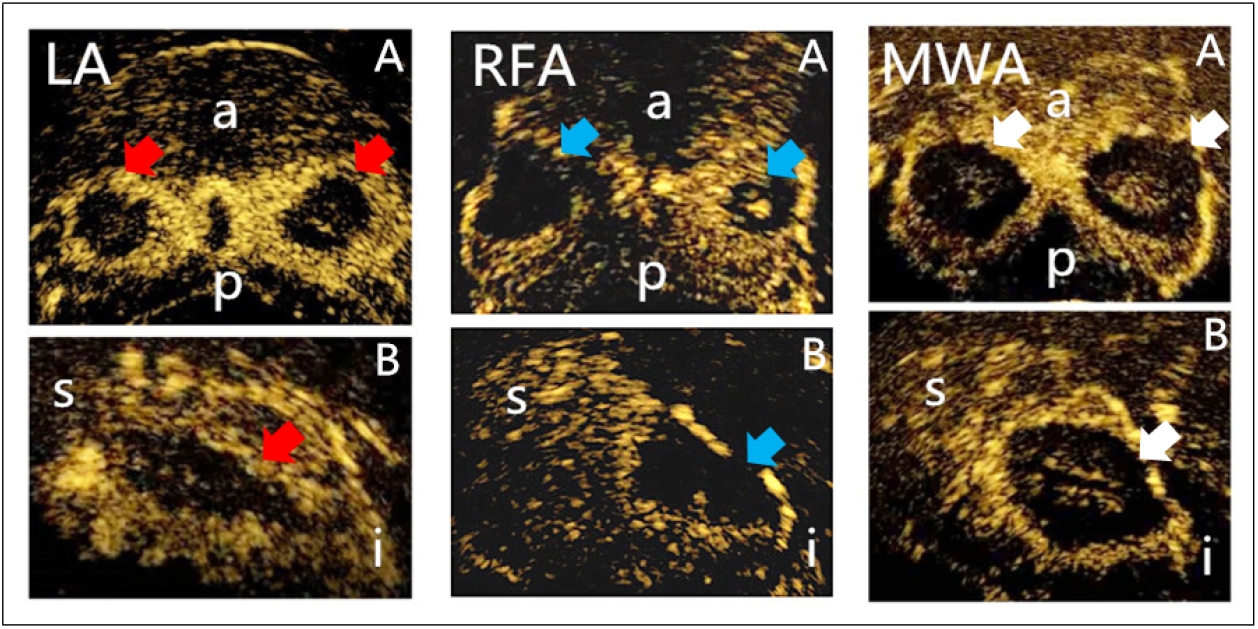
**The transverse (A) and longitudinal (B) section images after LA, RFA, and MWA were observed in CEUS** (red arrow, LA lesion; blue arrow, RFA lesion; white arrow, MWA lesion; a, anterior; p, posterior; s, superior; i, inferior).

Using the clinically commonly used doses, the R values were 0.89±0.02, 0.72±0.01, and 0.65±0.03 for FRA, MWA, and LA, respectively, which were significantly different (*P* =0.027). The size of lesion underwent MWA (30 w/120 s) was the highest (2.17±0.10 ml), followed by LA (3 w/1200 J; 1.51±0.20 ml), and RFA (30 w/120 s; 0.79±0.07 ml). It was revealed that size of lesion was also significantly different among the three techniques (*P* =0.001) (Table 1).

**Table 1.**
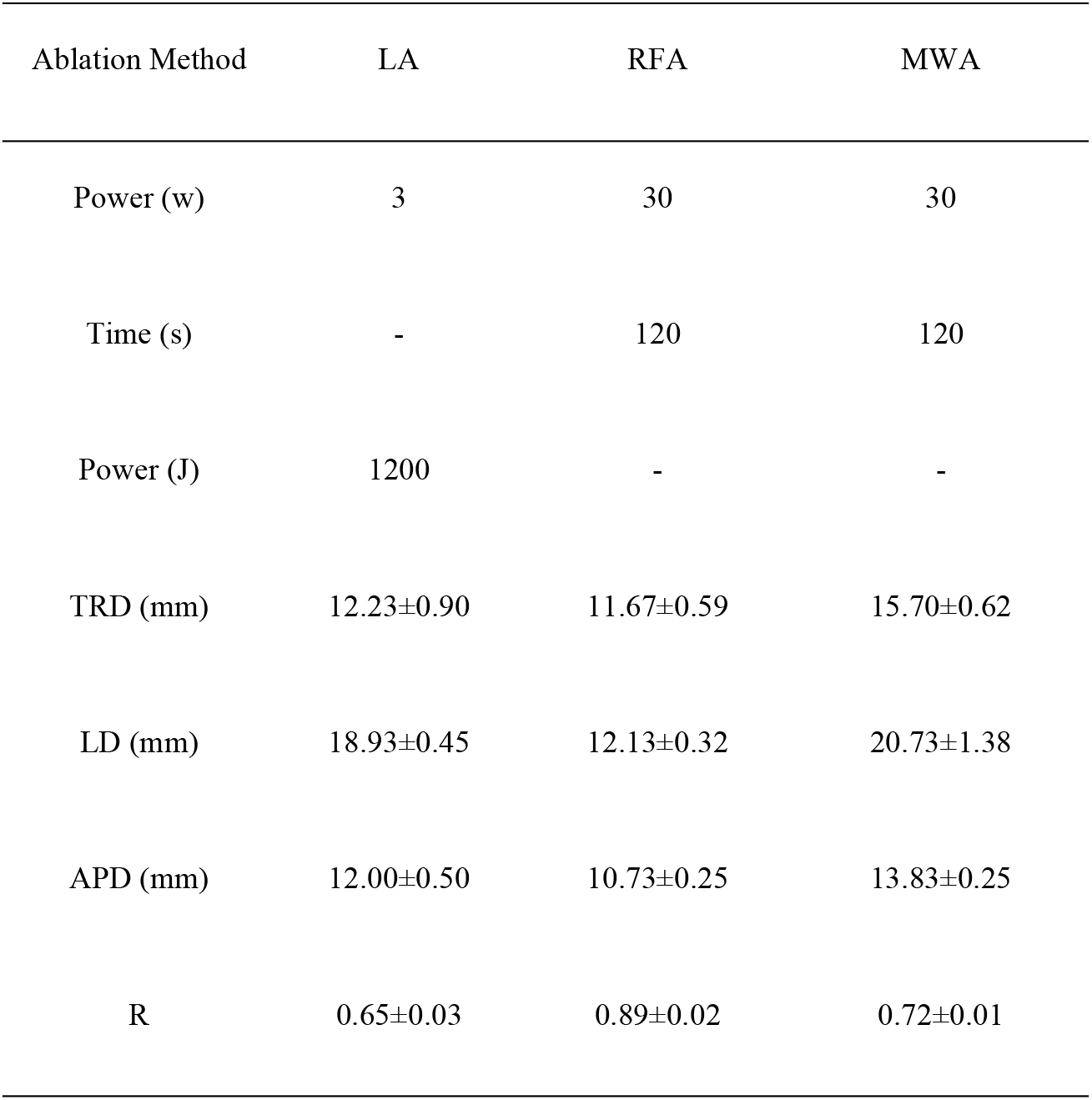
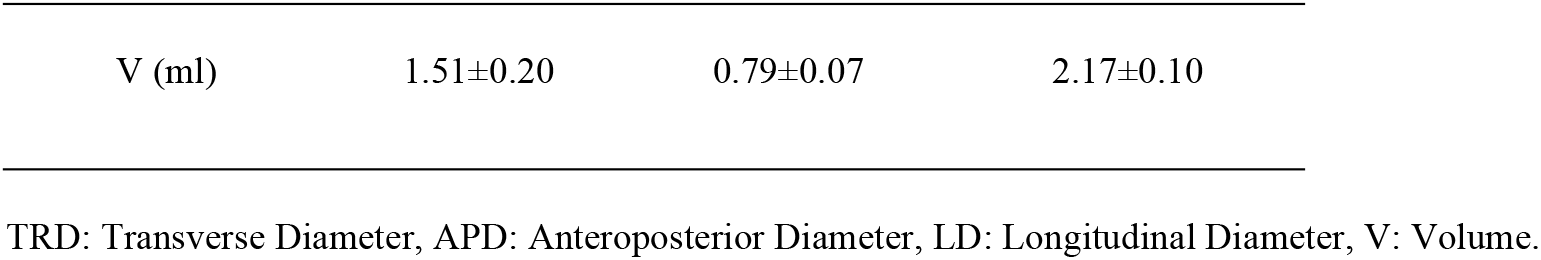
Comparing length, size, and shape (R) of lesions underwent LA, RFA, and MWA.

| Ablation Method | LA | RFA | MWA |
| --- | --- | --- | --- |
| Power (w) | 3 | 30 | 30 |
| Time (s) | - | 120 | 120 |
| Power (J) | 1200 | - | - |
| TRD (mm) | $12.23\pm0.90$ | $11.67\pm0.59$ | $15.70\pm0.62$ |
| LD (mm) | $18.93\pm0.45$ | $12.13\pm0.32$ | $20.73\pm1.38$ |
| APD (mm) | $12.00\pm0.50$ | $10.73\pm0.25$ | $13.83\pm0.25$ |
| R | $0.65\pm0.03$ | $0.89\pm0.02$ | $0.72\pm0.01$ |
| V (ml) | 1.51±0.20 | 0.79±0.07 | 2.17±0.10 |

TRD: Transverse Diameter, APD: Anteroposterior Diameter, LD: Longitudinal Diameter, V: Volume.

## Discussion

LA has been applied in clinical practices for the treatment of prostate disease, and its satisfactory effectiveness has been previously reported [9,18]. The light is transmitted by the optical fibers to the tissues, generating heat, and high temperature induces irreversible damages to tissues [19]. It is noteworthy that RFA and MWA utilize high-frequency current and electromagnetic wave, respectively, to induce the high-speed vibration and friction of ions and polar molecules in local tissues, which in turn generate high temperature and induce necrosis of tissues and cells [20]. These techniques were extensively used in clinical practices for the treatment of benign and malignant solid tumors of liver, lung, and thyroid[21–23], and the effectiveness and safety profiles of those techniques were documented [10,14]. However, a limited number of studies have employed these techniques for the treatment of prostate disease. Prostate is located at the center of pelvic cavity, surrounding the urethral canal, as well as being adjacent to various vital organs. For both BPH and localized PCa, complete ablation of the lesion, maintaining the patency of urethral canal and avoids damage to surrounding organs, is essential for local treatments. Therefore, “shape-appropriate” ablation for local prostate may contain positive therapeutic consequences for patients. In the current research, LA, RFA, and WWA with the clinically commonly used doses were undertaken for ablation of prostate in dogs [9,14,15]. The volumes and shapes of the lesions were compared to provide an evidence for the selection of “shape-appropriate” ablation techniques for the treatment of prostate diseases.

The findings of this study showed that LA, RFA, and MWA could be used for the local ablation of prostate. TRUS examination showed that hyperechoic regions were gradually expanded during LA, RFA, and MWA, which was followed by acoustic tail. These changes were caused by diffusion of bubbles of water content in tissues induced by high temperature. A number of studies reported application of the hyperechoic regions for distinguishing ablation areas. However, due to irregular shape, unclear boundaries, and influence of gas on acoustic wave transmission, the regions might be easily missed. In addition, the present study revealed that the hyperechoic region was transmitted to distant areas along the interstitial space. Therefore, further studies are required to verify whether the hyperechoic regions could be used to precisely distinguish the ablation areas.

Previous studies [10,24] demonstrated the feasibility of transrectal CEUS, and showed a high consistency between transrectal CEUS and pathological results of the necrotic area. In the current study, transrectal CEUS was used to assess the characteristics of lesions underwent ablation. The results demonstrated that the lesions were spherical or oval, without contrast agent filling. However, the boundaries of the lesions were clear, and the shapes of the lesions were regular, which were in agreement with previous findings [10,24]. The outcomes suggested that utilizing transrectal CEUS after prostate ablation could distinguish the ablation areas in dogs. However, further studies need to be conducted to indicate whether transrectal CEUS could precisely identify the ablation area in human. The radial lines of the lesions were measured, which showed that the shapes of the lesions using different ablation techniques had unique characteristics. The lesions were mainly spherical or oval on the transverse section of transrectal CEUS images. However, on the longitudinal section images, lesions underwent LA were “drop-shaped” shaped, while lesions received RA and MWA were more spherical. In addition, R value, reflecting changes in 3D shape varied as well. For instance, R values were 0.89±0.02, 0.72±0.01, and 0.65±0.03 for lesions underwent RFA, MWA, and LA, respectively. These findings demonstrated that 3D shapes of lesions received RFA and LA were more spherical and oval, respectively, and the shapes of lesions underwent MWA were between spherical and oval.

The present study also measured the sizes of the lesions received the three ablation techniques under clinically commonly used doses. The results uncovered that the size of MWA lesions was the highest, followed by lesions received LA and RFA. The ablation mechanisms and powers required were different among the three techniques; consequently, the results lack the comparability in certain degrees. However, for the three techniques used in the present study, the doses were all the clinically commonly used doses. We speculated that the evident variations in the size of lesions could be associated with the following causes: MWA is not influenced by current conduction or dryness and carbonization of tissues, thus, the size of lesion is large during a short-period time; however, in the RFA process, the temperature in the surrounding tissues reaches a certain height and carbonization occurs, hindering the radio frequency current propagation and limits the range of ablation.[25]. Although LA is not influenced by resistance; however, the temperature in the target point is relatively high, which induces the tissue carbonization and cavitation, and consequently influences the penetrating capacity of laser [26]. These findings were in agreement with the previous findings in ablation of thyroid tissues [13].

The results related to the sizes and shapes of the prostates of the dogs that underwent the three ablation techniques are of great significance for future application in clinical practice. The lesions underwent RFA had the lowest size, and their shape was more spherical, demonstrating that the mentioned method could be used for treatment of lesions with a relatively small size, or tumors surrounded by vital organs, such as urethral canal, rectum, and seminal vesicle. The lesions underwent MWA had the highest size, while their shape was spherical, which could be advantageous for ablation of tumors with a relatively large size. The size of lesions received LA was between those underwent RFA and MWA, and their shape was more oval, suggesting that the proposed method is highly appropriate for ablation of BPH. As lesions showed a “narrow-stripe shape” [27], the proposed method could possibly reduce urethra distortion, and could be therefore used for ablation of “stripe-shaped prostate cancer” close to the urethral canal. However, further studies are required to indicate whether these techniques could be effectively helpful in clinical practice.

There were several limitations in the present study. Firstly, only the clinically commonly used doses were taken in this study into account to explore the size and shape of lesions, while the other parameters were not investigated. Secondly, only prostate of normal dogs were ablated in this study, while the ablation area in tumor tissues or human prostate might be different. Finally, for the same ablation technique, the characteristics of lesions might vary due to the use of different types of ablation devices. Further researches need to be conducted to indicate whether the findings could be applied in clinical practice.

